# Exploring the Influence of Pore Shape on Conductance and Permeation

**DOI:** 10.1101/2024.04.18.589791

**Authors:** David Seiferth, Philip C. Biggin

**Affiliations:** Clarendon Laboratory, Department of Physics, University of Oxford, Oxford, OX1 3PU, UK; Structural Bioinformatics and Computational Biochemistry, Department of Biochemistry, University of Oxford, Oxford, OX1 3QU, UK

## Abstract

There are increasing numbers of ion channel structures featuring heteromeric subunit assembly, exemplified by synaptic α1β_B_ Glycine and α4β2 Nicotinic receptors. These structures exhibit inherent pore asymmetry, but the relevance of this to function is unknown. Furthermore, molecular dynamics simulations performed on symmetrical homomeric channels often leads to thermal distortion whereby conformations of the resulting ensemble are also asymmetrical. When functionally annotating ion channels, researchers often rely on minimal constrictions determined via radius-profile calculations performed with computer programs, such as HOLE or CHAP, coupled with an assessment of pore hydrophobicity. However, such tools typically employ spherical probe particles, limiting their ability to accurately capture pore asymmetry. Here, we introduce an algorithm that employs ellipsoidal probe particles, enabling a more comprehensive representation of the pore geometry. Our analysis reveals that the use of non-spherical ellipsoids for pore characterization, provides a more accurate and easily interpretable depiction of conductance. To quantify the implications of pore asymmetry on conductance, we systematically investigated carbon nanotubes (CNTs) with varying degrees of pore asymmetry as model systems. The conductance through these channels shows surprising effects that would otherwise not be predicted with spherical probes. The results have broad implications not only for the functional annotation of biological ion channels, but also for the design of synthetic channel systems for use in areas such as water filtration. Furthermore, we make use of the more accurate characterization of channel pores to refine a physical conductance model to obtain a heuristic estimate for single channel conductance. The code is freely available, obtainable as pip-installable python package and provided as a webservice.

## Introduction

Advances in structural biology, coupled with the emergence of artificial intelligence-driven structure prediction algorithms, have significantly expanded our knowledge of three-dimensional ion channel and nanopore structures in various conformational states. Notably, AlphaFold(1) and cryo-electron microscopy have played a pivotal role in unravelling the intricate details of these molecular structures. However, despite these remarkable strides, the functional annotation of these structures remains a critical challenge. To address this gap, several tools have been developed, such as HOLE(2), Caver(3), Moleonline(4) and the Channel Annotation Package (CHAP)(5), which enable the analysis of the physical dimensions and characteristics of the pores running through ion channels.

One of the first programs to compute the pore radius of an ion channel was HOLE (2). The HOLE program utilizes a Monte Carlo simulated annealing procedure to determine the optimal route for a sphere with varying radius to traverse through the channel On the other hand, CHAP(5), combines calculations of pore radius, hydrophobicity, and water density to predict hydrophobic gates in ion channels. CHAP is written in C++ and utilizes the trajectory analysis framework of the MD simulation software GROMACS.

Other tools, such as MOLEonline(4) and CAVER(3), have also been employed for the detection and characterization of channels, pores, and tunnels within biomacromolecules and do not use a probe based algorithm for path finding. MOLEonline provides information about channel-lining residues, physicochemical properties, and identifies cavities using Voronoi diagrams and molecular surfaces. CAVER employs a grid-based approach and convex approximation to determine the lowest-cost centreline path between a given starting point and the molecule’s surface(6). In later versions, CAVER also uses Voroni diagrams(7). The visualization of the CAVER or MOLEonline path, despite not being probe-based in its algorithm, relies on the utilization of probe particles. By utilizing the maximum radii for each node along the path, researchers can effectively visualize and analyse the characteristics of the identified channels or pores. Although some methods offer the chance to visualize detailed (highly asymmetric) surfaces, probe-based methods offer a more intuitive way to interpret the geometry in terms of simple statistics like radii.

The HOLE program, in particular, has been commonly used by structural biologists as a way to quickly (and robustly) provide key information on pore geometry. Until recently, the vast majority of transmembrane channel structures were highly symmetrical, usually as a direct consequence of homomeric subunits forming higher-order multimer structures coupled with symmetrical averaging techniques in the structural processing. However, as structural biology techniques improve, the number of structures that exhibit inherent asymmetry is increasing and understanding its potential role and relationship to function is gaining attention. Minniberger *et al*.(8) explored the asymmetry and ion selectivity properties of bacterial channel NaK mutants derived from ionotropic glutamate receptors. They observed structural asymmetry in the upper part of the selectivity filter, proposing that local asymmetrical conformational changes could efficiently alter ion channel function without requiring large-scale conformational transitions. Similarly, *Roy et al*.(9) studied a nonselective sodium-potassium ion channel with structural asymmetry in the selectivity filter, suggesting that different degrees of conformational plasticity are required for conducting potassium and sodium.

Zhang *et al*.(10) proposed a model for asymmetric activation of the 5-HT_3_A serotonin receptor, while Shi *et al*.(11) highlighted the permeation mechanisms of sodium (Na^+^) and potassium (K^+^) in NaK channels. Lewis *et al*.(12) observed structural changes between Na^+^ and K^+^ -bound states, accompanied by increased structural heterogeneity in Na^+^. Chugunov *et al*.(13) reported asymmetric gating at the selectivity filter in molecular dynamics simulations of the TRPV1 pore upper gate. These studies collectively emphasize that conformational plasticity and structural asymmetry in the selectivity filter play crucial roles in regulating ion selectivity and conductance. Furthermore, local asymmetrical conformational changes emerge as an energetically efficient means to modulate ion channel function without requiring global conformational transitions across all subunits.

The presence of asymmetry in ion channel pores brings into question as to whether the use of a spherical probe is appropriate. In fact, this was anticipated many years ago, and the HOLE software already provides a so-called CAPSULE option to measure pore anisotropy with a spherocylinder. However, an asymmetrical pore can be more accurately described with an ellipsoidal probe. Along the channel axis, two radii are reported corresponding to the longer and shorter axes of the ellipsoid. The volume of the pore characterised with ellipsoidal probe particles will be larger than the volume found when using spherical probe particles. To that end and building upon the well-established HOLE program and its MDAnalysis Python API (14) we have developed a program which we call PoreAnalyser. It is available as a pip-installable python package (https://github.com/bigginlab/PoreAnalyser) and interactive web-service (https://poreanalyser.bioch.ox.ac.uk) and allows users to capture asymmetry via the use of an ellipsoidal probe. To demonstrate its use, we report here an assessment of the influence of asymmetry on conductance predictions.

To systematically quantify the implications of pore asymmetry, we investigated Carbon Nanotubes (CNTs) as model systems, exploring the conductance through channels with varying degrees of pore asymmetry. The investigation builds upon the cylindrical approximation model, a fundamental physical model employed to predict channel conductance. Hille(15) characterized homogeneous conducting materials by a bulk property known as resistivity *ρ*_*bulk*_, representing the inverse of conductivity *κ*_*bulk*_. The conductance model, treating the channel as a cylinder, considered both pore resistance *R*_*pore*_ associated with the length of the pore and the so-called access resistance at the pore entrance (*R*_*access*_) and as in equation 1.

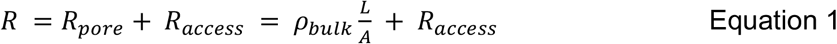

where *A* denotes the cross-sectional area and *L* the length of the cylinder. Subsequent studies, such as that by Smart *et al*. (16), utilized Hille’s cylindrical model but neglected access resistance, approximating an ion channel as a series of cylinders with different radii as determined by HOLE analysis. Mejri *et al*. (17) extended this work using CNTs by revealing that increasing the CNT diameter enhanced conductance values, while an increase in tube length decreased conductance, suggesting a series-like action of access resistance and pore resistance. Manghi *et al*.(18) further investigated the relative contributions of channel and access resistances in experimentally measured currents.

Carbon Nanotubes (CNTs) are established model systems for ion channels in the realm of molecular dynamics (MD) studies(19,20). Their controllable dimensions make them invaluable tools for investigating ion transport phenomena. CNTs have been explored in the context of de-wetting processes within confined hydrophobic regions of channels. Confined hydrophobic regions in ion channels and CNTs may undergo transitions between wet and dry states to gate the pore closed without physical constriction of the permeation pathway – a phenomenon called hydrophobic gating(21). Beckstein *et al*.(22) contributed to the understanding of de-wetting mechanisms by proposing a hydrophobic gating mechanism for nanopores, emphasizing the role of nanopore geometry, diameter and the hydrophobicity of pore lining residues. In the context of this current study, we focused on channels wide enough to rule out de-wetting transitions - we only consider conductive CNTs with radii larger than 4 Å whose pores are consistently wetted.

Mendonça *et al*.(23,24) delved into the intricacies of water diffusion within deformed CNTs characterized by varying degrees of eccentricity. Their investigations provided insights into the behavior of water molecules within these nanoscale conduits, particularly emphasizing the freezing of water under certain conditions. Ezaee *et al*.(25) studied monoatomic fluid flow through elliptical carbon nanotubes, systematically examining the influence of tube size, temperature, and pressure gradient on the fluid dynamics. Balme *et al*.(26) contributed to the understanding of ionic transport through hydrophobic nanopores, observing modifications in the ion solvation structure when nanopore diameters were less than 2 nm and varying NaCl concentrations. Additionally, Mejri *et al*.(17) focused on the role of water models (SPC/E, TIP3P, TIP4P/2005) in the study of ionic conductance using molecular dynamics simulations. Their findings indicated that the choice of water model had no significant impact on the conductance trend concerning increasing tube diameter; however, it did influence conductance values. Notably, hydrogen bond analyses revealed similar water arrangements within the pore for both the SPC/E and TIP4P/2005 models, highlighting the robustness of certain water models in capturing essential aspects of ion transport through CNTs. Mejri et al. studied the impact of different chemical CNT functionalization and geometric parameters such as radius and chirality on conductance through MD simulations with an external potential(27). Together, these studies contribute to an understanding of the complex interplay between CNT characteristics, water behavior and ionic conductance.

We thus chose CNTs as models systems where we can easily control the extent of pore-asymmetry in the system. The conductance through these CNTs show surprising effects that would otherwise not be predicted with spherical probes. Building on the cylindrical approximation, we propose an improved physical model to predict conductance. The results have broad implications not only for the functional annotation of biological ion channels, but also for the design of synthetic channel systems for use in areas such as water filtration. We anticipate that PoreAnalyser can more readily take into account asymmetrical properties of ion channel structures.

## Methods

### Pore geometry calculations

We align the principal axis of the input structure to the z-axis by default, although this can be modified as per user preference (users of the web-interface can simply upload their own coordinates). To calculate the pore geometry, we employ probe-based pathway finding using spherical probe particles as a first step and then we transform each sphere into an ellipsoid. Initially, HOLE is executed using a spherical probe particle, with the centre of mass (COM) as the default starting point. The position of the probe is optimized in subsequent parallel planes to maximize its radius without overlapping the van der Waals sphere of any ion channel atom. **Fig. 1** illustrates the schematic representation of van der Waals spheres, the positions of their centres in the xz-plane of a channel aligned to the z-axis, and a surface representation of the pore. HOLE is integrated into our Python workflow with its MDAnalysis Python API(14)

**Figure 1.**
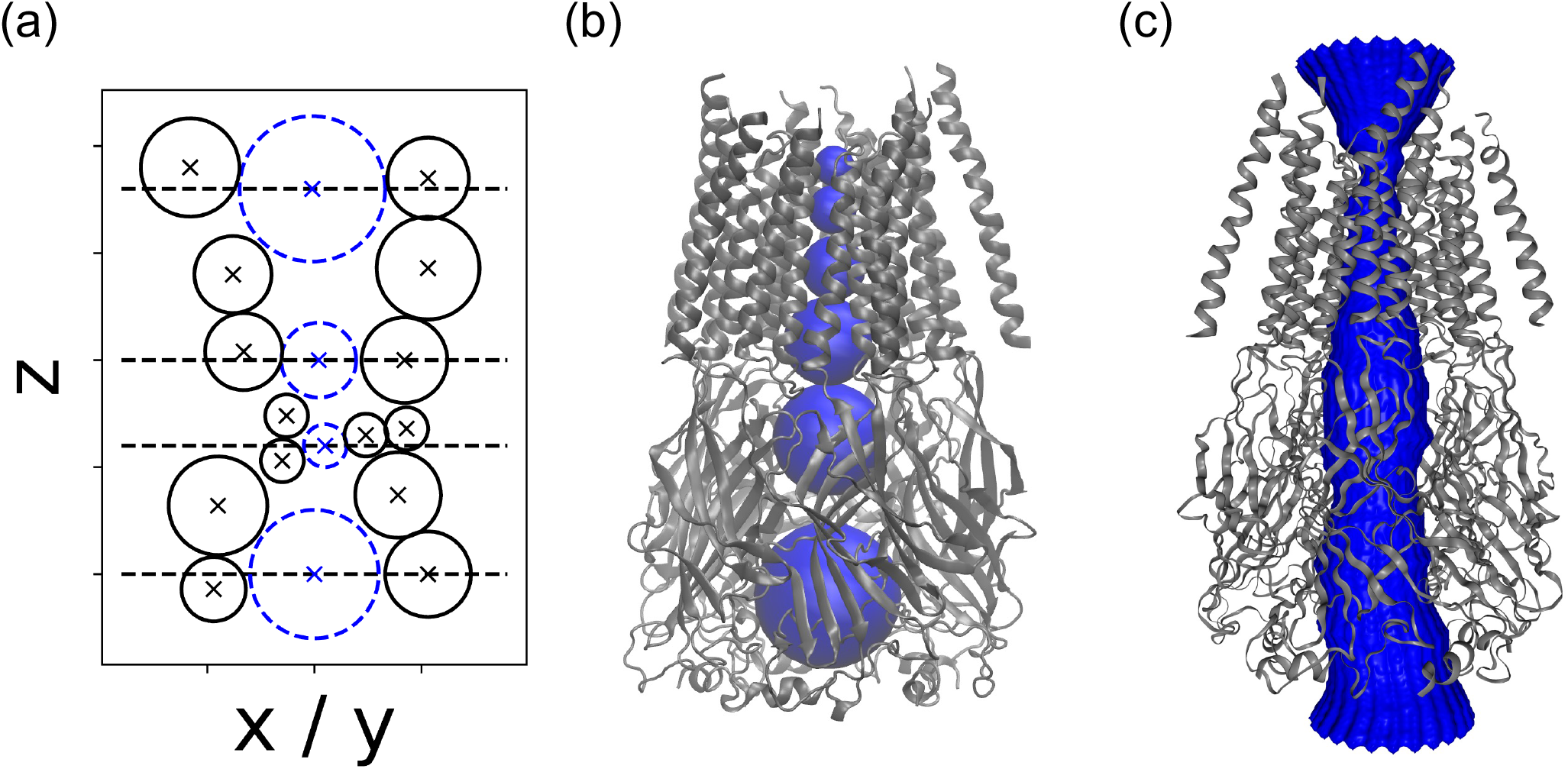
Pore geometry calculations with spherical probe particles used in HOLE (2) or CHAP(5). **(a)** The position of a spherical probe is iteratively adjusted in parallel planes to achieve the maximum radius while ensuring there is no overlap with the van der Waals sphere of any atom within the ion channel. **(b)** Positions of a few probe particles along the pathway of the pore within the α1β_B_ Glycine receptor(28) (PDB 7tvi) **(c)** Surface representation of the pore of the α1β_B_ Glycine receptor.

To incorporate an ellipsoidal probe particle, we iteratively transform the spherical probe particles into ellipsoids. To initiate the calculation, we load the output file from HOLE containing the positions and radii of the probes. Then, we perform a loop through all the spherical probe particles and initialize an ellipsoid using the parameters obtained from the HOLE output. The parameters of the ellipsoid include the position of its center [*x, y, z*], orientation *θ*, and radii [*a, b*], where *a ≥ b*. The smaller radius, *b*, remains constant along the z-coordinate. To optimize the parameters, we employ a Nelder-Mead four-dimensional optimization algorithm, first using smaller bounds for the parameters [*x, y, a, θ*], and then with larger boundaries to further increase the ellipsoid (**Fig. 2**). Finally, we compare the volumes of the pores based on the use of spherical and ellipsoidal probe particles. This comparison provides insights into the impact of probe shape on pore geometry calculations.

**Figure 2:**
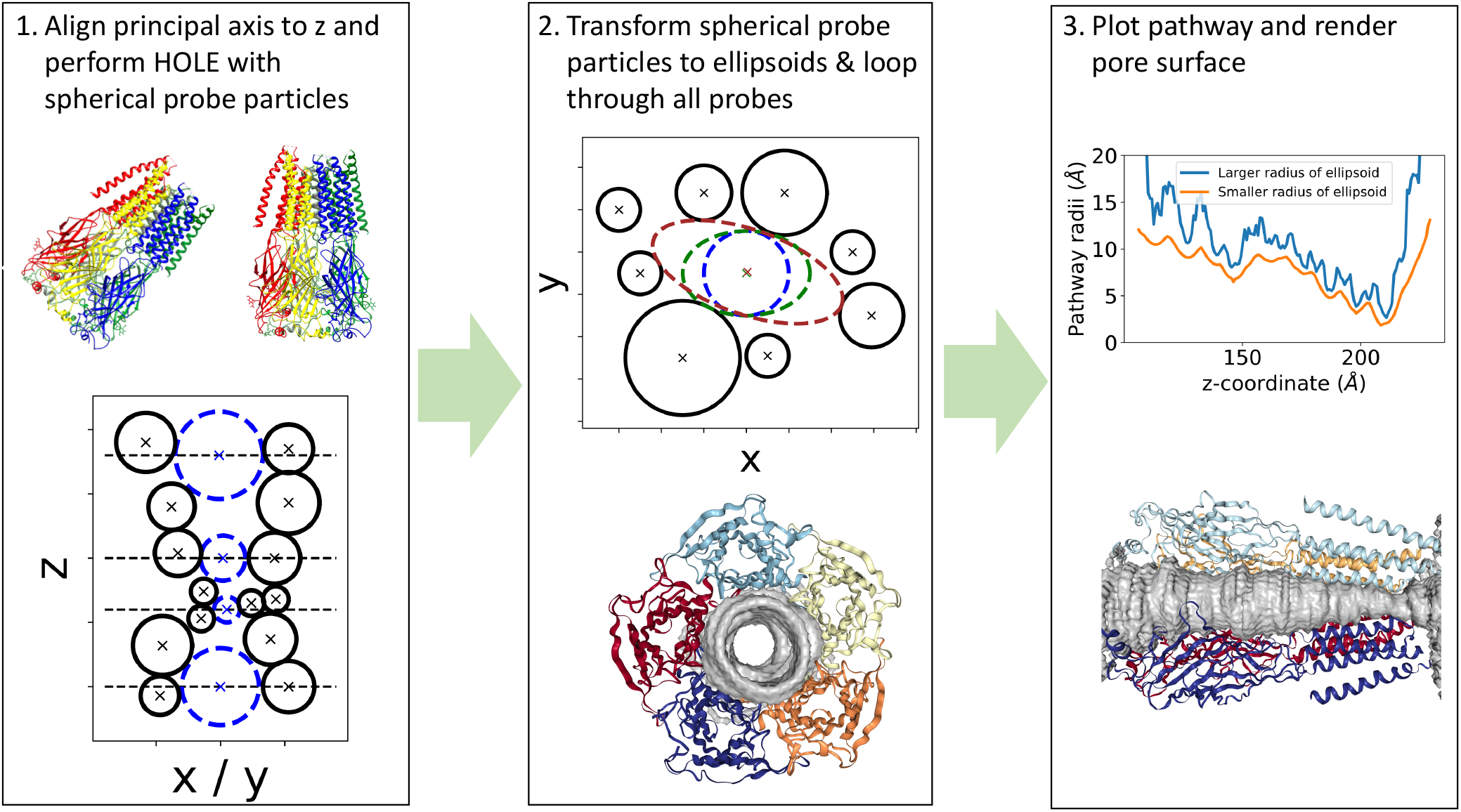
Workflow diagram of the web-service and the python package. The input protein is aligned to the z-axis and then a HOLE analysis is performed (1). The HOLE spheres are then iteratively transformed into ellipsoid (2). As a final step, the pathway is plotted and the pore surface visualized (3).

### Model systems

We employ pristine armchair carbon nanotubes (CNTs) capped with hydrogen atoms as a model system, featuring a length of 50 Å, a dimension that enables them to span a lipid bilayer completely. The key characteristic of the CNTs lies in the variation of their radius, carefully selected to be wide enough to prevent de-wetting of the pore, ensuring stable hydration throughout the simulation duration. All CNT atoms have zero partial charge. The simulation setup involves the insertion of the CNT into a bilayer (see **Fig. 3**) composed of 128 1-palmitoyl-2-oleoyl-sn-glycero-3-phosphocholine (POPC) lipids, facilitated by the InflateGro method(29). The system is solvated with a 0.15 M KCl solution. Atomistic molecular dynamics simulations are conducted using GROMACS(30), wherein the CNT is modelled using the OPLS all-atom force field(31), united-atom lipids, and the SPC/E water model(32). Simulation parameters include a time integration leap-frog with a 2 femtosecond time step, constraints applied to bonds using the LINCS(33,34) algorithm, and the particle mesh Ewald method(35,36) employed for handling long-range electrostatic interactions. The temperature is maintained at 300 K. To preserve orientation, the z-coordinates of the CNT are restrained, with harmonic restraints of 500 kJ/mol/nm^2. Introducing varying degrees of asymmetry, we manipulate the area in the xy-plane *A*_*xy*_ = *πr*^*2*^ = *π a b*, where *r* represents the radius of the CNT circular plane, *a* denotes the larger, and *b* the smaller radius of the ellipsoid. The asymmetry is quantified by the ratio of the smaller to the larger elliptic radius (½, ¾, and 1), corresponding to different degrees of eccentricity 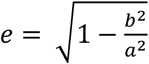, depicted in **Fig. 3 (a)-(c)**. The cross-sectional area *A*_*xy*_ is preserved for varying degrees of eccentricity. To retain a non-circular shape, harmonic restraints are defined such that a major axis of *2a* and a minor axis of *2b* are enforced. Notably, the pores remain stably hydrated throughout the entirety of the simulations, providing a robust foundation for investigating the influence of varying degrees of asymmetry on the behavior of CNTs within lipid bilayers. We study (8,8), (10,10), (12,12), (14,14), (16,16) and (18,18) pristine armchair CNTs with different degrees of eccentricity, where the native (8,8) and the (14,14) have HOLE radii of 3.4 Å and 7.0 Å respectively (see **Table 1**).

**Table 1.** Carbon nanotubes (CNTs) of length 50 Å and varying HOLE radii.

| Armchair CNT of type (n,m) | (8,8) | (10,10) | (12,12) | (14,14) | (16,16) | (18,18) |
| --- | --- | --- | --- | --- | --- | --- |
| HOLE radius (Å) | 3.4 | 4.7 | 5.9 | 7.0 | 8.3 | 9.8 |

**Figure 3.**
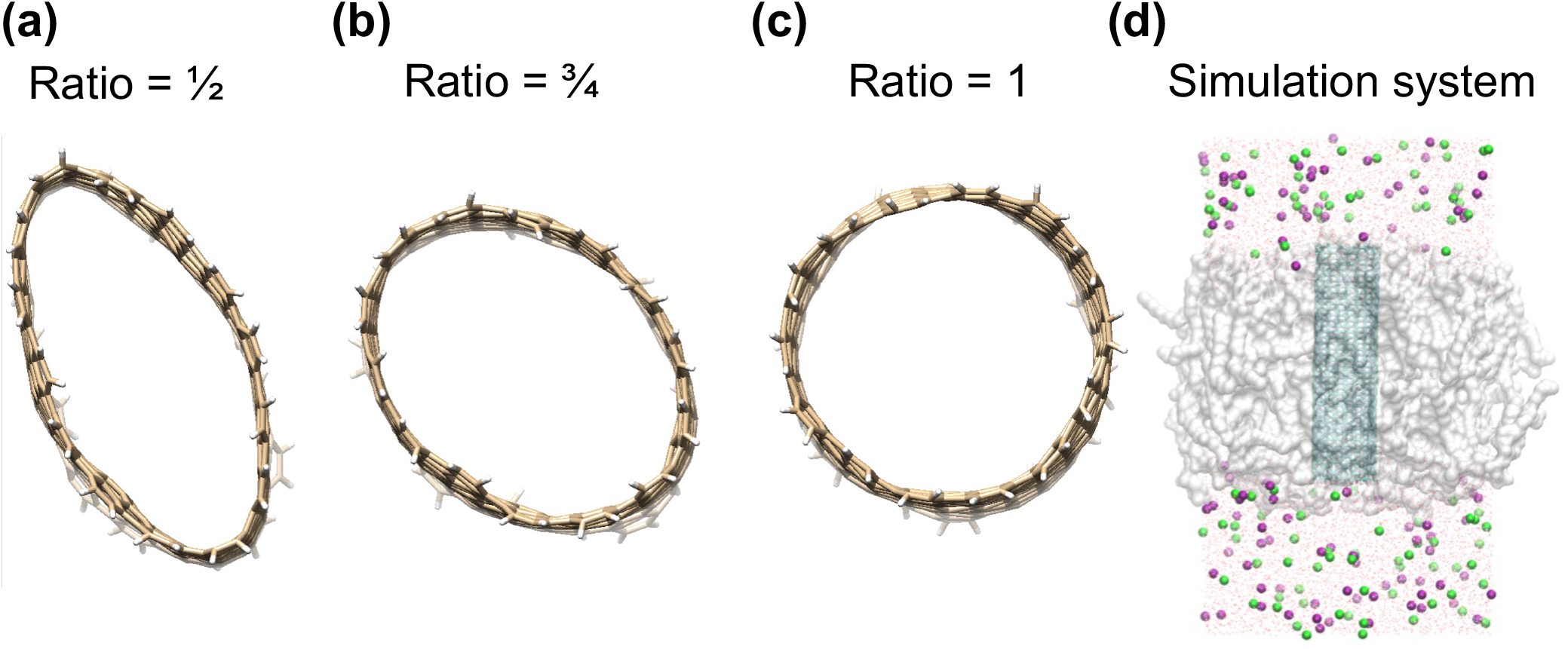
Cylindrical carbon nanotubes with varying degrees of asymmetry. View of xy-plane. **(a)** Elliptical xy-plane with ratio of ½ between smaller and larger radius. **(b)** elliptical xy-plane with ratio of ¾ between smaller and larger radius. **(c)** Circular xy-plane (smaller and larger radius of ellipse are identical). The area A_xy_ = πab in the xy-plane is conserved and virtual bonds are defined to maintain the degree of asymmetry. **(d)** Cartoon of the simulation box with the carbon nanotube in cyan, the lipid in grey translucent spheres, water in red and white wire and K^+^ and Cl^-^ ions.

We denote a (14,14) armchair CNT with length 50 Å as 14 x 50 in this article. The CNT type, denoted by the pair of integers (n, m), signifies that a specific CNT can be derived by rolling a graphene sheet characterised by lattice vectors 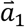 and 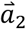. In this configuration, atoms align theoretically atop one another when positioned at a distance of 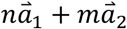 on the graphene sheet. Ion conduction was measured in 250 ns simulations at 150 mM KCl concentration and in the presence of a +500mV and -500mV transmembrane potential difference. This was applied by imposing an external, uniform electric field in the membrane normal direction. Umbrella sampling was performed to obtain one-dimensional single-ion potential of mean force profiles for potassium and chloride passing through the pore of different CNTs with varying degree of asymmetry. Simulation protocols were similar to those detailed above for equilibrium MD simulations. The distance between the respective ion and the centre of mass of the CNT structure along the z-axis of the simulation box was chosen as the collective variable. Starting configurations were generated by swapping positions of the respective ion with water molecules along the permeation pathway. The minimum of the biasing potential was shifted by 0.2 nm between each umbrella sampling window. A force constant of 500 kJ/mol/nm^2^ restrained the z-coordinate of sodium to the respective distance to the centre of mass of the protein. Unbiasing was performed with the weighted histogram analysis method (WHAM) using the gromacs implementation(37).

## Results and discussion

### A simple case study with a biological ion channel

To demonstrate how the geometrical profile of the pore is influenced by the use of an ellipsoidal probe, we investigated the pore geometry of the full-length zebrafish α1β glycine receptor (GlyR) as a case study. GlyR plays a crucial role in inhibitory neuronal signaling in the spinal cord and brainstem. Synaptic GlyRs are a heteromeric assembly of α and β subunits and hence asymmetric(28). We analyzed a cryo-EM structure of the full-length zebrafish α1βGlyR in glycine-bound state (PDB code 7TVI)*(28)*. Our analysis, as depicted in **Fig. 4**, reveals that the local minima of the pathway radii obtained using a spherical probe particle, representing the smaller radius, coincided with the local minima obtained from the larger radius of the ellipsoid. This observation indicates the consistency and reliability of our calculations. Moreover, we found that the ratio of pore volume based on the ellipsoidal and spherical probe particles was approximately 1.6. This indicates that using a spherical probe particle alone would lead to a significant underestimation of the actual pore volume. The incorporation of an ellipsoidal probe particle provides a more accurate representation of the true pore volume, offering valuable insights into the size and dimensions of the GlyR pore. These findings highlight the importance of considering different probe shapes when calculating pore geometry and emphasize the limitations of relying solely on spherical probe particles for accurate measurements.

**Figure 4.**
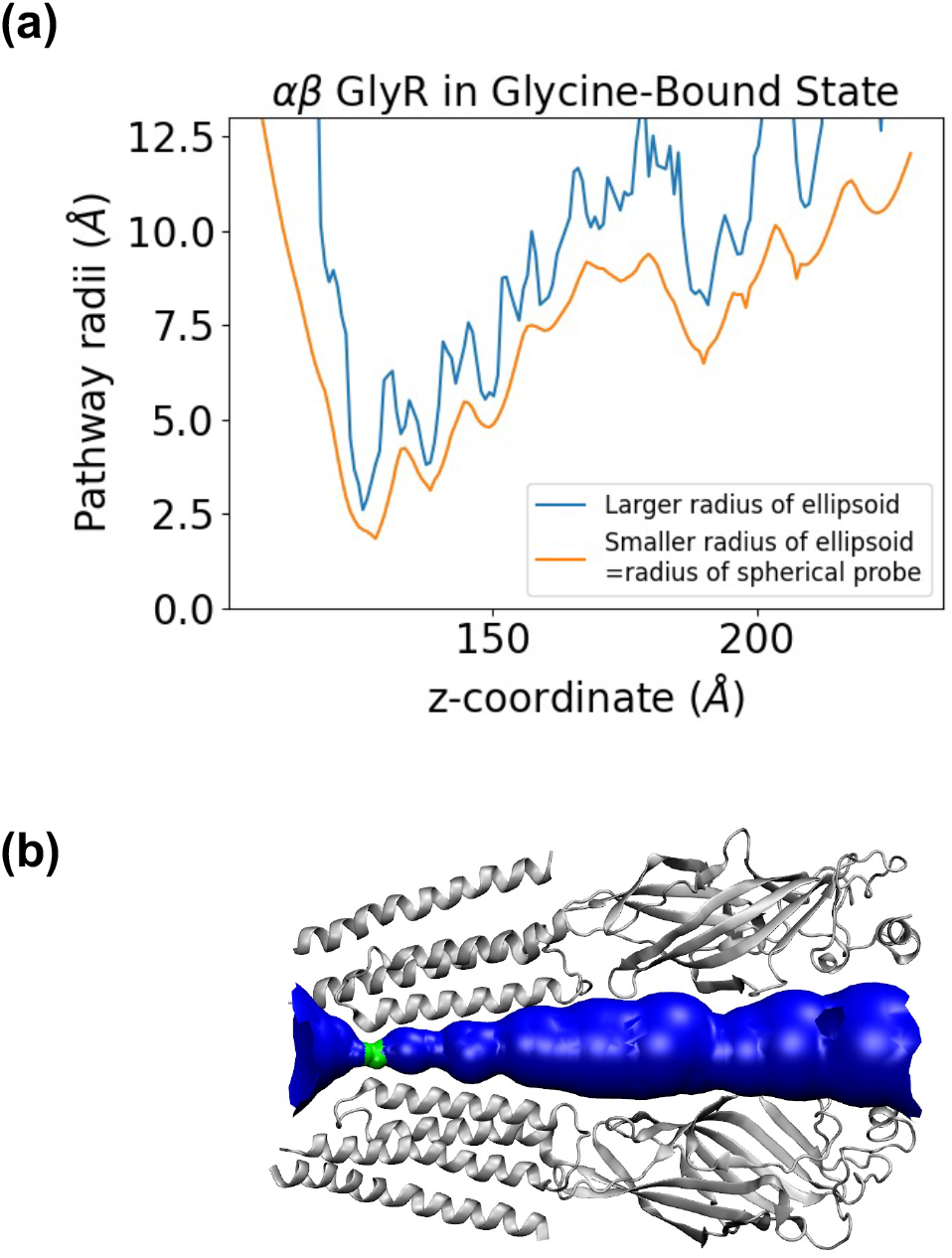
Case Study of the α1β_B_ GlyR Channel. **(a)** Pathway analysis for a cryo-EM structure of full-length zebrafish α1β_B_ GlyR with heteromeric subunit assembly in the presence of glycine(28) **(b)** cartoon showing the pore profile.

### Comparison with HOLE capsule option

As mentioned earlier, Smart *et al*.(16) introduced the CAPSULE option within the HOLE algorithm to allow the user to capture pore asymmetry, a feature that has yet to gain widespread adoption within the scientific community. Departing from the traditional approach of optimizing a single sphere, HOLE with CAPSULE option extends its capability to a spherocylinder, essentially an extension of a sphere. The spherocylinder is uniquely characterized by two independently moving centres, described by three properties: its radius *r* and the positions of these two centres with distance *L* between them(16). Central to the algorithm is the measurement of anisotropy, achieved by ensuring that the axis of the capsule aligns perpendicular to the channel direction vector. Unlike the original routine, which optimized the sphere’s radius, the CAPSULE option focuses on maximizing the area of the capsule on a given plane. Subsequently, this area is converted into an effective radius 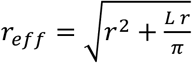, providing a parameter for ease of comparisons. In **Fig. 5**, we compare the performance of the HOLE CAPSULE algorithm with our PoreAnalyser package, where ellipsoidal probe particles are employed to comprehensively explore the intricacies of pore geometry, providing valuable insights into the relative strengths and limitations of these two distinct algorithms. The smaller elliptical radius *b* of the PoreAnalyser algorithm and the spherical radius *r* of HOLE are identical by construction and are plotted in blue in **Fig. 5**. The larger elliptical radius *b* of the PoreAnalyser algorithm is consistently larger than the effective radius *r*_*eff*_ of the capsule both for an elliptical CNT 10 x 50 model system with ratio of elliptical radii b/a = 1/2 depicted in **Fig. 5 (a)-(b)** and for a GlyR ion channel depicted in **Fig. 5 (c)-(d)**. The PoreAnalyser algorithm can accurately capture both axis of ellipsoidal probe particles whereas the HOLE capsule option underestimates the cross-sectional area. For the GlyR ion channel, the minima of larger and smaller radii of the PoreAnalyser algorithm coincide. When moving away from the narrowest constrictions, the differences between the major and the minor elliptical axis become more pronounced. The pore volume obtained from PoreAnalyser is significantly larger than the one obtained from HOLE capsule. For both systems, the PoreAnalyser algorithm describes the respective pore more accurately.

**Figure 5.**
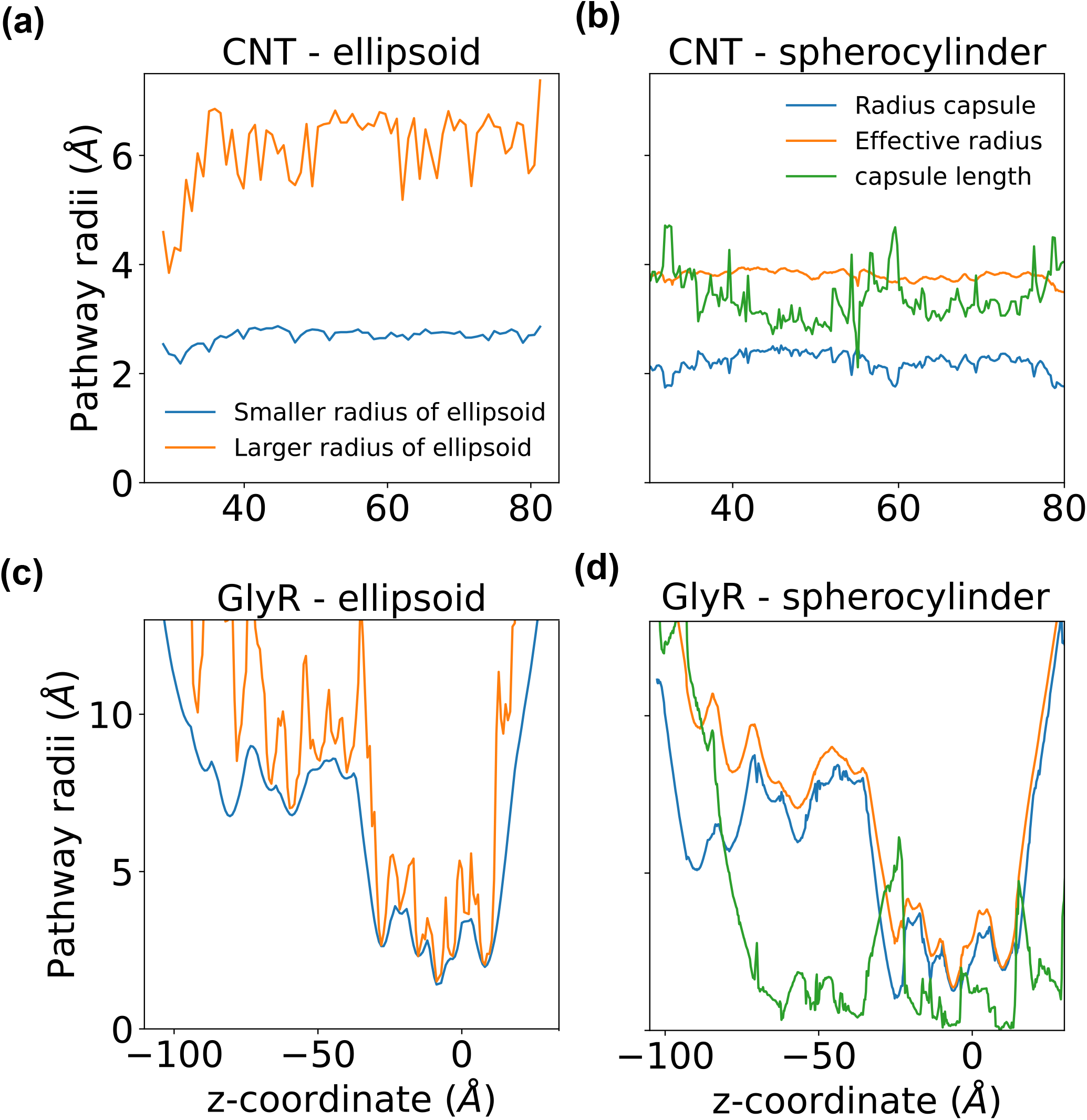
Comparison of pathways found with elliptic probe particle of PoreAnalyser with the CAPSULE option from HOLE. A carbon nanotube profile of a 10 x 50 system with ratio 1/2 obtained from PoreAnalyser **(a)** and HOLE CAPSULE option **(b)**. Profiles obtained from PoreAnalyser **(c)** and HOLE CAPSULE **(d)** for the Glycine receptor (PDB code 7TU9).

**Figure 6.**
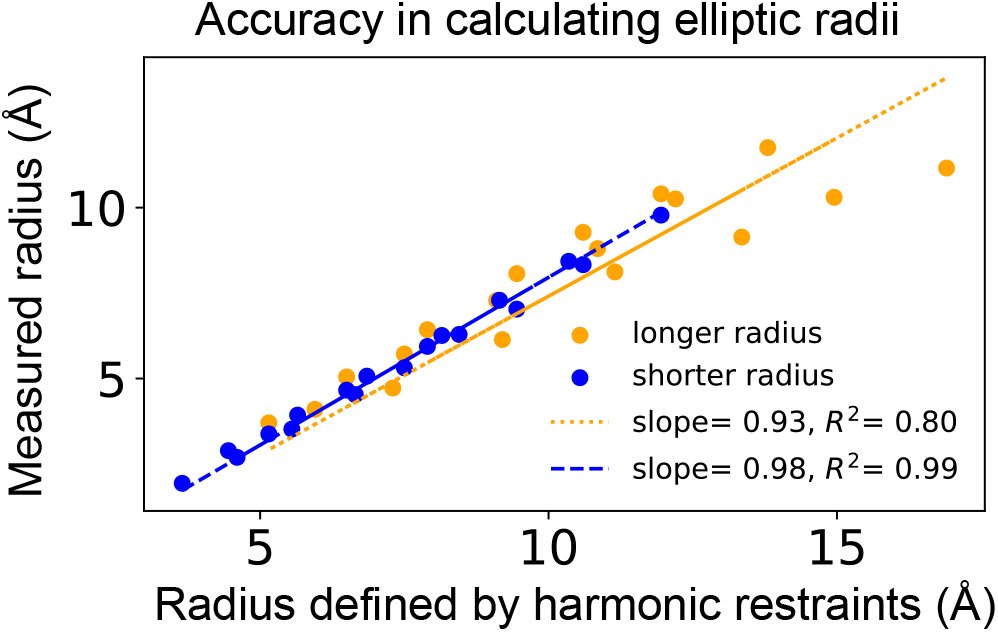
Evaluating the accuracy in calculating elliptic radii in 18 different carbon nanotubes. It can be seen that the longer radius is less well maintained than the shorter radius, particularly at radii above 9 Å.

### Carbon nanotubes to investigate conductance

To evaluate the accuracy of the PoreAnalyser algorithm when employing ellipsoidal probe particles, a comprehensive analysis was conducted on various Carbon Nanotube (CNT) model systems. The extent of ellipticity is denoted by two radii; *a* (longer radius) and *b* (shorter radius) for each CNT. We enforce the ellipticity through the use of harmonic restraints to maintain the elliptical shape with prescribed ratios *b/a* of 1, ¾, and ½ in **Fig. 4**. We thus first checked to see how well the restraints maintained the desired radius (**Fig. 4**). Notably, higher standard deviations were observed for CNTs with larger elliptical radii, signifying a pronounced variability when measuring the major axis. The pores identified using spherical probe particles effectively characterized the minor axis, offering a benchmark for comparison. We examined 18 distinct CNTs, each with six different areas in the xy-plane and three varying ratios between the longer and shorter radii (1, ¾, ½). The HOLE radii of the circular CNTs (corresponding to a ratio of radii of one) are listed in **Table 1**. As expected, the HOLE radius, indicative of the shorter radius, aligned well with the minor elliptical axis. However, it was observed that the larger radius was less accurately maintained (**Fig. 4**) as might be anticipated from the use of harmonic restraints in this way. Nevertheless, this level of accuracy was deemed acceptable.

Having satisfied ourselves the model systems are behaving as expected, we then used these CNTs with varying degrees of pore asymmetry as model systems to investigate how pore asymmetry affects ionic conductance. Smart *et al*.(16) predict the conductance *g* of a channel based on the pore geometry found by HOLE as

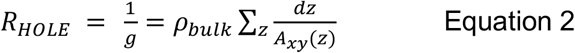

Where the resistivity *ρ*_*bulk*_ of the permeant ions is regarded as a function of the molar ion concentration *c*. The pore is approximated as a sequence of cylinders with area *A*_*xy*_(*z*) in the xy-plane and length *dz*. This model predicts higher resistance *R* (and lower conductance *g*) for longer and narrower pores. However, the model would predict the same conductance for a channel with circular *A*_*xy*_ = *πr*^*2*^ and for a channel with elliptical area *A*_*xy*_ = *πab* if the cross-sectional area *A*_*xy*_ is the same. We measured the conductance and the barrier to permeation for potassium and chloride in four different CNT systems with varying degree of asymmetry. As one would expect, a higher barrier to permeation corresponds to a lower conductance (**Fig. 7**). Here, the conductance for potassium is lower than for chloride and consistent with this trend, the energetic barrier is higher for potassium than for chloride. We study four different CNT systems with increasing cross-sectional area (CNT 8 x 50 has the lowest cross-sectional area and CNT 14 x 50 the highest), and the conductance increases with increasing cross-sectional area *A*_*xy*_ in line with Equation 2. When we vary the ratio of the elliptical axes while preserving the cross-sectional area *A*_*xy*_, we find that a system with ratio 1, corresponding to a spherical cross-sectional area, has consistently higher conductance values than the same system with ratio ½, corresponding to major axis twice as long as the minor axis. The energetic barrier decreases for less elliptical shapes.

**Figure 7.**
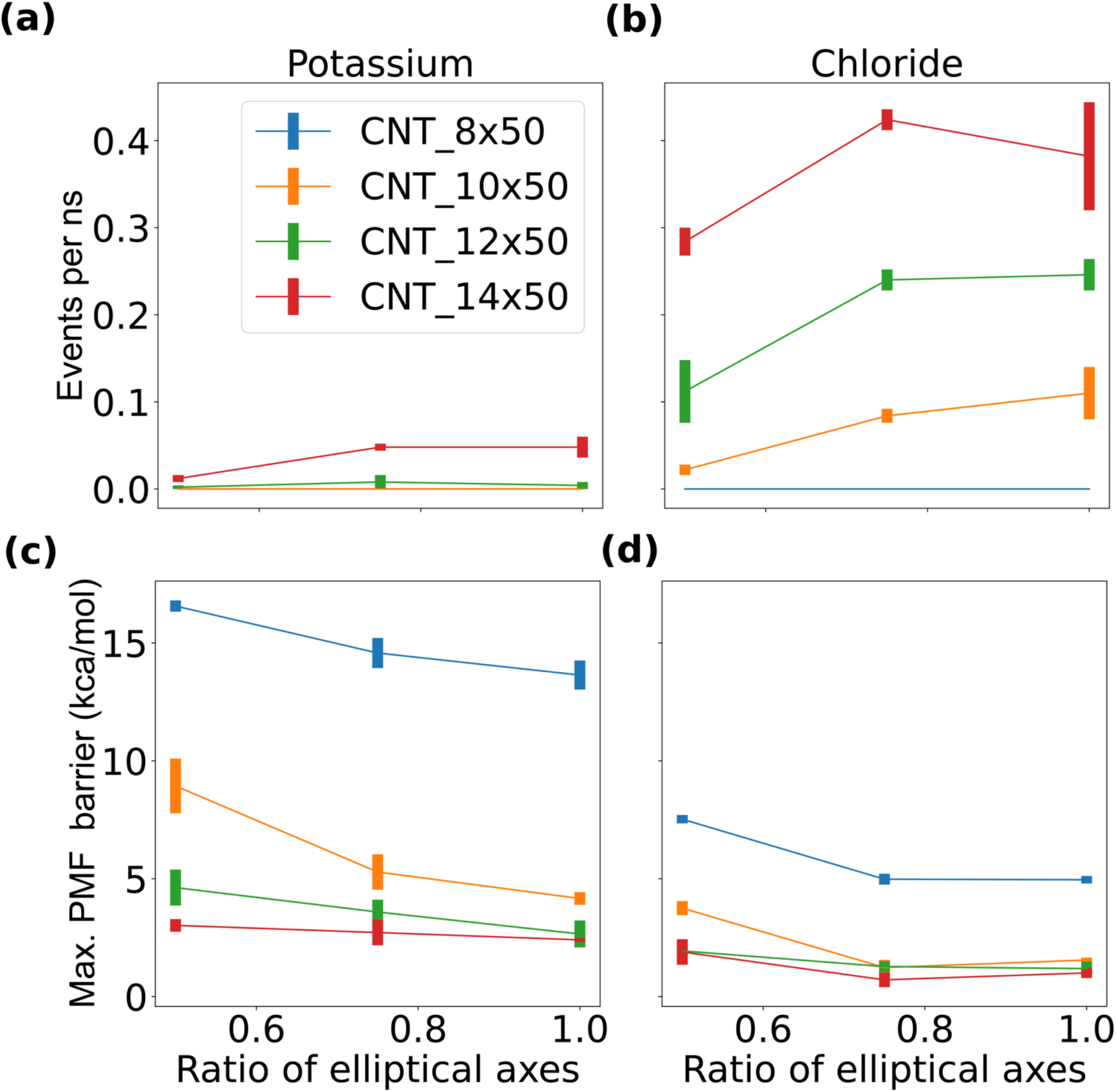
Barrier for ion permeation and ion conductance for carbon nanotubes (CNTs) as a function of the ratio between of the elliptic radii. The conductance is measured as events per ns observed in multiple 250 ns repeats with external potential of 500 mV for **(a)** potassium ions and **(b)** chloride ions. The energetic barrier is the maximum of the Potential of Mean Force (PMF) obtained from umbrella sampling for **(c)** potassium ions and **(d)** chloride ions.

We do not observe potassium or chloride conductance events in the repeats of 250 ns in the narrowest CNT system (CNT 8 x 50). The energetic barrier for potassium and chloride to permeate the pore increases with higher eccentricity of the cross-sectional area. The differences are more pronounced for narrower CNTs with smaller cross-sectional areas (**Fig. 7**). We next calculated the potential of mean force (PMF)s and estimated the conductance for the ion parameters obtained from Li *et al*.(38). The barrier for ion permeation and the ion conductance are depicted in **SI Fig. S1**. In a recent study we found that ion parameters can influence free energy barriers significantly(39) but when comparing **Fig. 7**. and **SI Fig. S1**, we see a similar trend for energetic barriers and conductances.

We can obtain a measure of the hydration shell along the reaction coordinate, by simply reporting the mean and standard deviation of how many water molecules are within 3 Å of the respective ion (**Fig. 8 c, d**). The hydration shell of potassium is distorted in the CNT with ratio ½ between minor and major axis of the elliptical cross-sectional area. The radial distribution function (RDF) for potassium (**Fig. 8e**) and chloride (**Fig. 8f**) in bulk and in the narrow CNT system reveals that for potassium, the second peak, corresponding to the second hydration shell, is significantly reduced compared to bulk. Potassium inside a CNT with ratio ½ between minor and major axis has a smaller RDF compared to potassium inside a spherical CNT (ratio 1).

**Figure 8.**
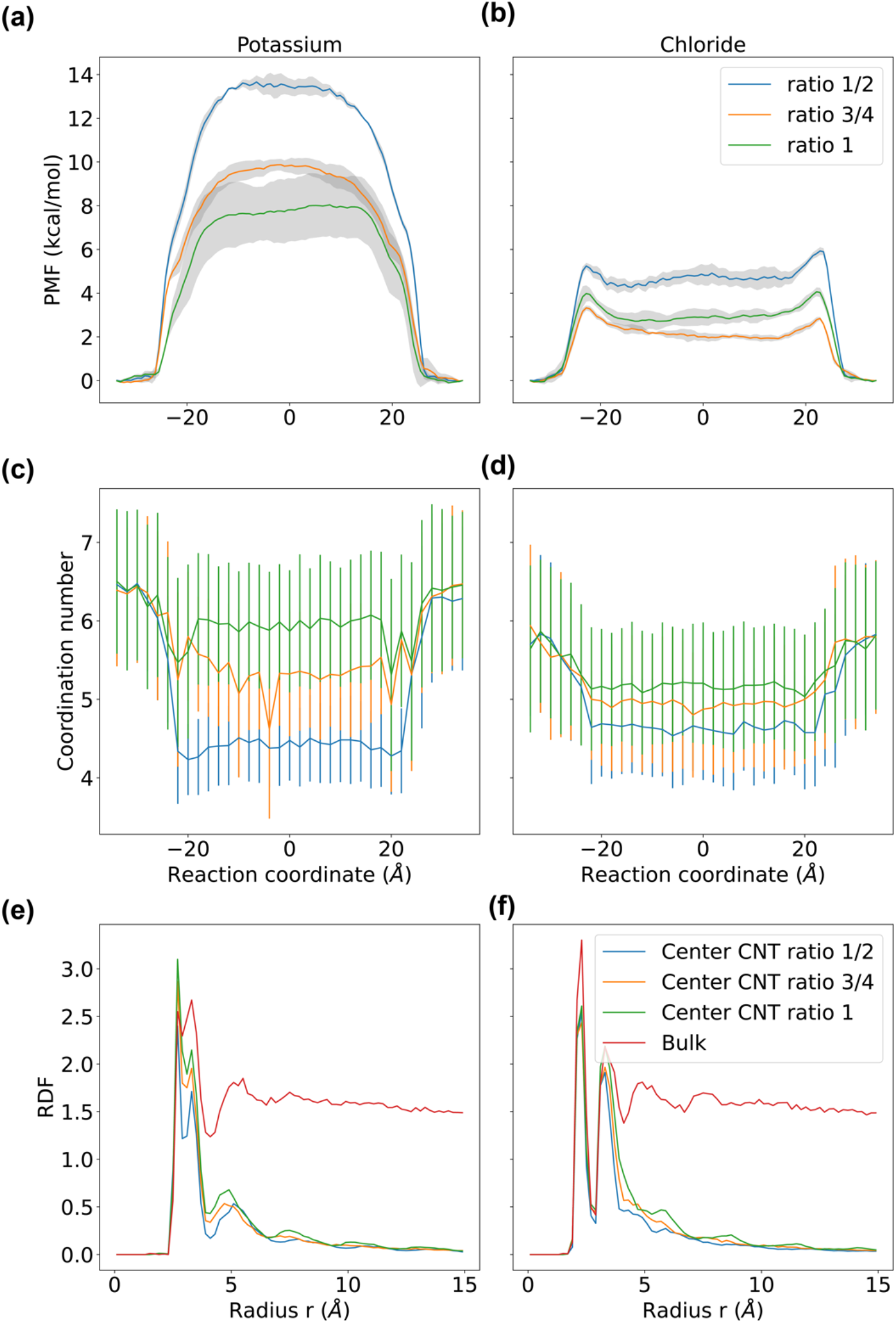
Potential of mean force (PMF). PMFs for Potassium **(a)** and chloride **(b)** to permeate 8 x 50 carbon nanotubes (CNTs) with different degrees of asymmetry. The reaction coordinate is parallel to the z-axis of the simulation box and the distance to the z-coordinate of the center of mass of the CNT. The hydration shell is plotted as a function of the reaction coordinate for potassium **(c)** and chloride **(d)** for a narrow CNT. The radial distribution function (RDF) of potassium **(e)** and chloride **(f)** in bulk and in the center of a 8 x 50 CNT with varying degree of asymmetry.

### Physical model to predict conductance

In the development of a physical model to predict conductance through ion channels, Hille’s approach^14^ initially considered a cylindrical approximation with length *L* and cross-sectional area *A*, allowing the resistance *R* to be expressed as:

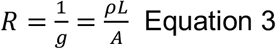

where *ρ* represents bulk resistivity. This simplistic model was subsequently refined (2,40) suggesting a more accurate representation of ion channels as a series of stacked cylinders, where the resistance accumulates. Considering ohmic principles and utilizing the HOLE software, which measures cross-sectional areas *A(z)* along the channel axis *z*, the refined resistance model becomes:

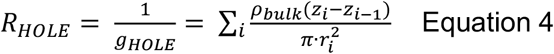

where the resistivity *ρ* is approximated with bulk property *ρ*_*bulk*_ and *r*_*i*_ denotes the radius of a spherical probe particle. However, relying on bulk property resistivity *ρ* = 1/*κ* becomes problematic, as conductivity *κ* depends on the diffusion coefficients of ions. The bulk conductivity *κ*_*bulk*_ of a KCl solution with concentration *c* is defined as:

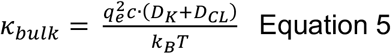

where *q*_*e*_ is the elementary charge, *D*_*κ*_ and *D*_*CL*_ are diffusion coefficients of potassium and chloride ions, *κ*_*B*_ is the Boltzmann constant, and *T* is temperature(26). To refine the model for ion channel conductance further, we introduce a conductivity model, expressing the conductivity *κ*(*a*_*i*_, *b*_*i*_) as a function of the radii *a*_*i*_ and *b*_*i*_ of ellipsoidal probe particles. For larger radii, the ion movement is relatively unconstrained, resulting in *κ*(*a*_*i*_, *b*_*i*_) ≈ *κ*_*bulk*_, while narrower constrictions with smaller radii lead to reduced conductivity *κ*(*a*_*i*_, *b*_*i*_) < *κ*_*bulk*_. Hence, we can further adapt the model for channel resistance / conductance based on the PoreAnalyser profile to:

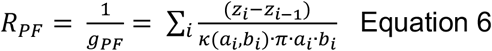

The conductivity function is represented by a double-sigmoid function with parameters *c*_1_ and *c*_2_, given by:

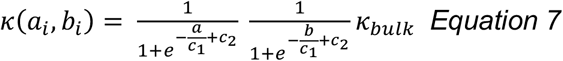

To fit parameters *c*_1_ and *c*_*2*_ of the model, training data from 18 Carbon Nanotube (CNT) systems, encompassing 6 CNTs with distinct cross-sectional areas *A*_*xy*_ and three different elliptic radii ratios (*a* and *b*), are utilized. Every model system of the training data corresponds to a point in **Fig. 9a**. In **Fig. 9b**, the ratio *κ*(*a*_*i*_, *b*_*i*_)/*κ*_*bulk*_ is illustrated as a function of varying radii, keeping one radius constant while the other varies. The conductance of these model systems is calculated based on their mean radii and compared to Molecular Dynamics (MD) conductance. The conductivity model

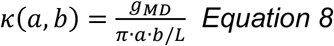

is employed for fitting, and **Fig. 9a** depicts the ratio *κ*(*a*_*i*_, *b*_*i*_)/*κ*_*bulk*_ in a two-dimensional plot as a function of both radii, with training data points plotted in the plane spanned by the radii *a* and *b*. When utilizing the bulk conductivity *κ*_*bulk*_ to predict the conductance of CNTs with eq (5), an overestimation of conductance is evident, as depicted in **Fig. 9c**. However, employing the conductivity model from equation (7) yields an improved correlation between MD conductance *g*_*MB*_ and the heuristic conductance *g*_*PF*_, resulting in an *R*^*2*^ value of 0.916.

**Figure 9.**
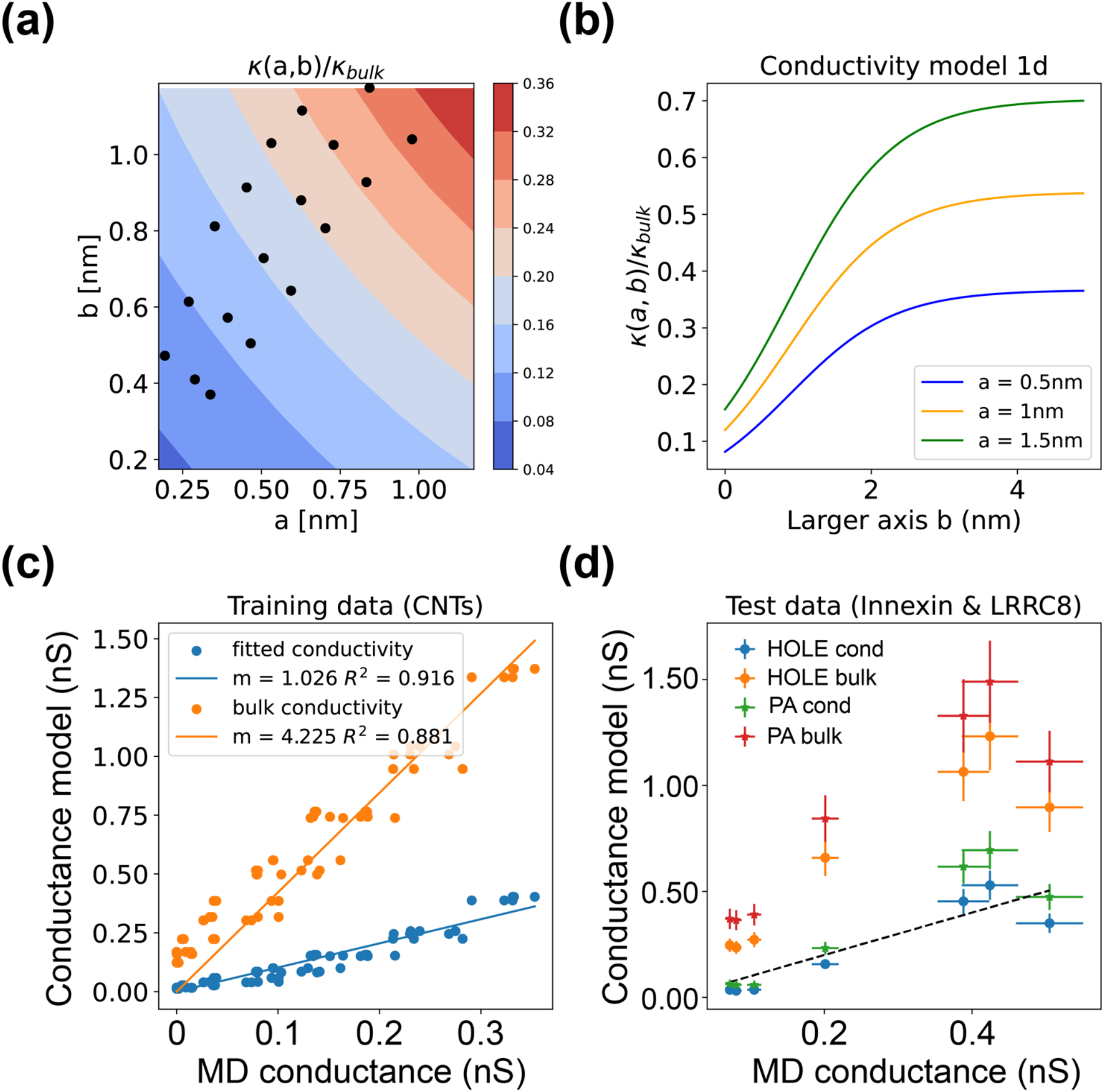
Conductivity model for KCl in confined environments characterized by radii a and b. The color surface of the ratio κ(a,b)/κ_bulk_ characterized by the two radii a and b **(a)**. One-dimensional sigmoid function with one radius fixed **(b)**. Comparison of MD conductance (considered to be ground truth) and conductance model using bulk conductivity and conductivity model **(c)**. Predicting conductance values for biological ion channels (see **Table. 2**) using the bulk conductivity or the conductivity model based on a HOLE (HOLE bulk or HOLE cond) or a PoreAnalyser (PA bulk or PA cond) profile **(d)**.

To further validate our enhanced physical conductance model, we extend the comparison to biological ion channels by embedding various Innexin and LRRC8 structures (see **Table 2**) in a POPC membrane and measuring MD conductance through simulations with an external potential. In **Fig. 9d**, we compare MD conductance and the heuristic based on HOLE or PoreAnalyser pathways, both with and without the conductivity model, for the Innexin(41,42) and LRRC8(43,44) test systems. The use of PoreAnalyser or HOLE pathway profiles in conjunction with bulk conductivity consistently overestimates conductance.

**Table 2.**
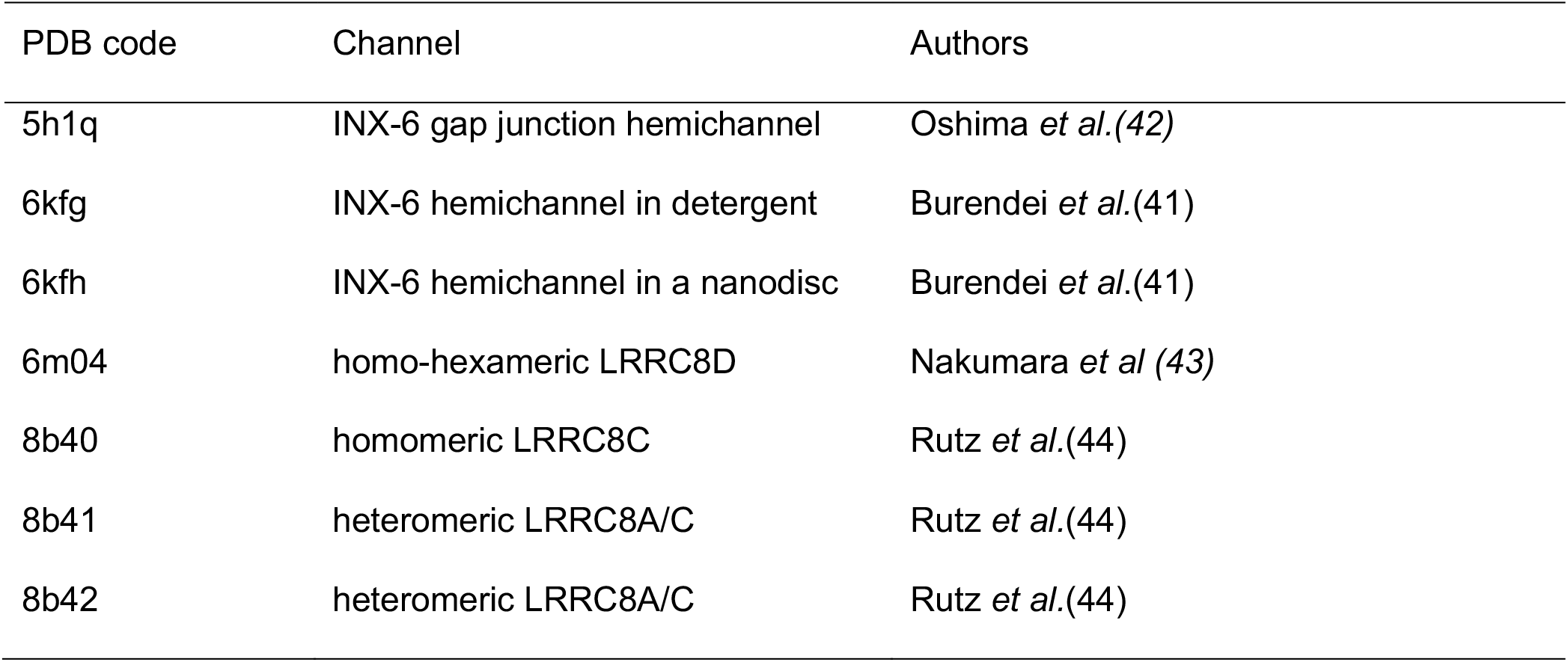
Innexin and LRRC8 structures used to test the conductance model.

Conversely, employing the conductivity model from eq. (5) with a HOLE pore profile often results in underestimation. The optimal agreement between MD conductance and the heuristic is achieved by using the conductivity model *κ*(*a, b*) in combination with the PoreAnalyser profile based on ellipsoidal probe particles. The errors of the MD conductance correspond to one standard deviation of three 250 ns simulations with external potential. To estimate the errors of the conductance model, we analyse the pore pathway for ten frames from the simulations and calculate the standard deviation.

To further enhance the understanding of the conductance model in equation (4) and highlight the contributions of different terms, we present the pore profile an innexin system in **Fig. 10a**. The conductivity along the channel axis is influenced by the radii *a* and *b* of the elliptical probe particle, as illustrated by the minima in **Fig. 10b**. The conductivity *κ*(*r*_*HOLE*_) of a HOLE profile is smaller than the conductivity *κ*(*a, b*) based on the PoreAnalyser profile. Ions, confined in a narrow environment, have a decreased mobility than ions in bulk solution. Calculating resistance based on the cylindrical approximation for each probe particle and plotting it along the channel axis (**Fig. 10c)**, we observe resistance maxima corresponding to the narrow constrictions in **Fig. 10a**. Substituting the bulk conductivity *κ*_*bulk*_ instead of the conductivity model *κ*(*a, b*) reduces channel resistance, resulting in an overestimation of channel conductance, as demonstrated in **Fig. 10c**.

**Figure 10.**
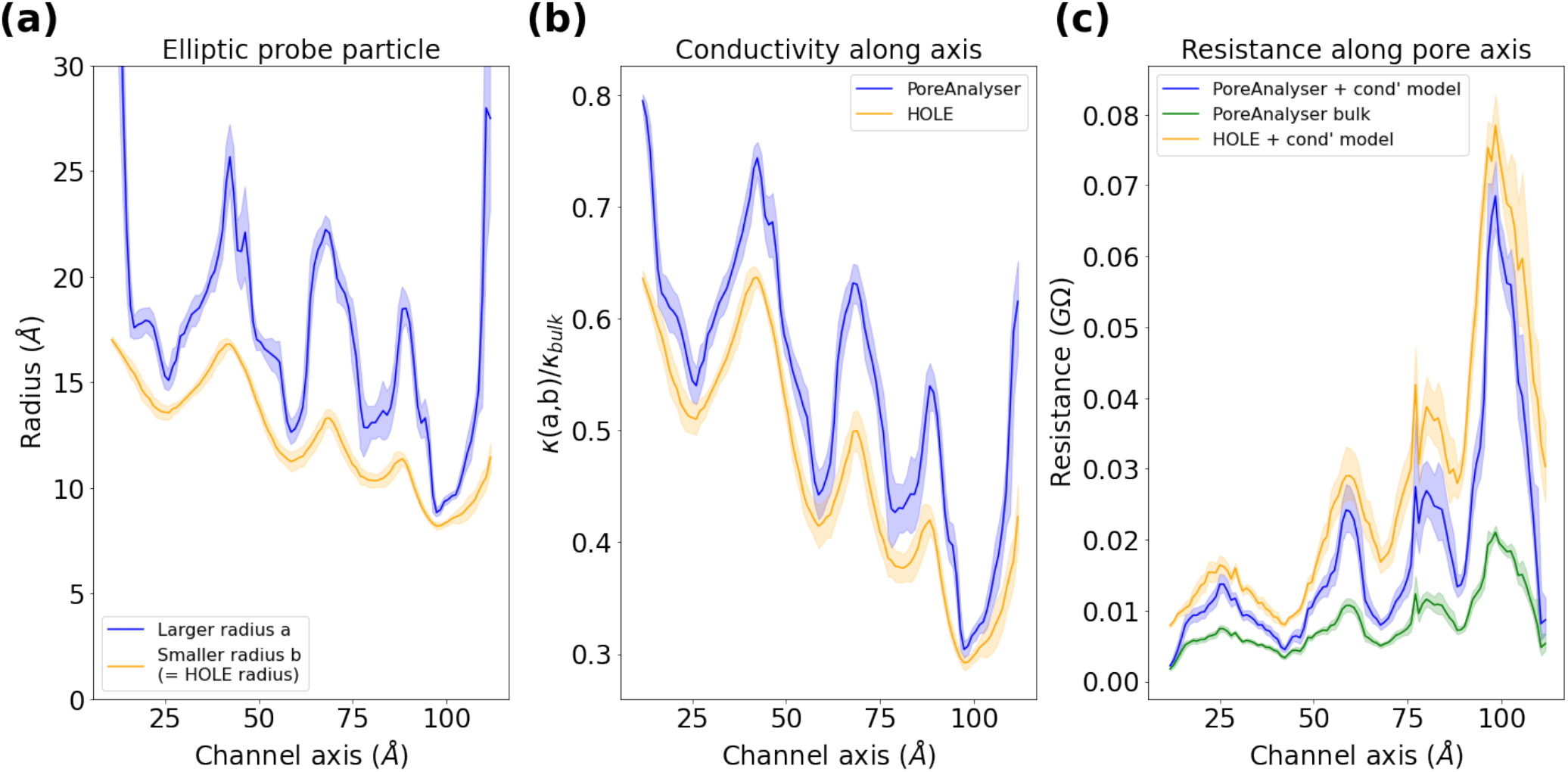
Conductivity model for Innexin (pdb code 6kfg, (41)). **(a)** Pore profile described by radii a and b along the channel axis. **(b)** Conductivity ratio κ(a, b)/κ_bulk_ along the channel axis. **(c)** Resistance along the channel axis employing pore profile based on HOLE and PoreAnalyser with and without the conductivity model (see equations 4 and 6). Errors are one standard deviation derived from an analysis of an ensemble of structures generated from a single MD trajectory.

## Conclusions

In this study, we focused on capturing pore asymmetry and its influence on conductance and barriers to permeation. Asymmetry can be observed in crystallographic or cryo-electron microscopy (cryoEM) structures, which can result from the heterogeneous composition of subunits within the channel complex. While crystal structures of channels with homogenous subunit decomposition exhibit a symmetrical arrangement, the symmetry is broken when these proteins are simulated in more realistic conditions. By incorporating our new features into the pore-analyser methodology, we were able to accurately capture and analyze the asymmetry of the pore, shedding light on its functional implications and providing a more comprehensive understanding of ion channel dynamics. The additional complexity of the pore computation adds a slight overhead to speed of the pathfinding process, which can take approximately 1 minute per frame when using 12 CPUs in parallel, but for most use-cases this is a reasonable timing. We demonstrated that consideration of asymmetry allows us to refine a physical conductance model to obtain a heuristic estimate for single channel conductance. This model allows users to predict single channel conductance without running costly simulations and can be used as an easy and cheap method for quickly predicting the functional state of new channel structures.

We were also keen to make the package as usable as possible and as such provide an interactive web-service (https://poreanalyser.bioch.ox.ac.uk) that allows users to calculate the pore profile of any input structure without the need for installation or downloads. Additionally, we have made the source code available on GitHub (https://github.com/bigginlab/PoreAnalyser or https://github.com/DSeiferth/PoreAn-alyser) and pip installable, which includes scripts for visualizing results in popular molecular graphics programs such as Chimera, VMD(45), and PyMOL(46). By providing these resources and making our software accessible online we aim to promote collaboration and further advancements in the field of pore analysis.

## Acknowledgements

DS is funded by the UKRI-BBSRC Interdisciplinary Bio-science Doctoral Training Partnership (BB/M011224/1). PCB acknowledge support from BBSRC (BB/S001247/1). Computing was supported via the Advanced Research Computing facility, Oxford, the ARCHER UK National Supercomputing Service and JADE (EP/T022205/1) granted via the High-End Computing Consortium for Biomolecular Simulation, (HECBioSim - http://www.hecbiosim.ac.uk), supported by EPSRC (EP/R029407/1).

## Data Availability

Copies of the source code can be obtained from https://github.com/bigginlab/PoreAnalyser and https://github.com/DSeiferth/PoreAnalyser and include scripts for visualizing results in the molecular graphics programs Chimera, VMD and PyMOL. The software is pip installable (pip install PoreAnalyser). The software can be used online without any installations via the webservice: https://poreanalyser.bi-och.ox.ac.uk.

## Author Contributions

DS and PCB conceived the project. DS developed the code. DS wrote the first draft and all authors contributed to revising and editing the manuscript.

