## Supplemental Information for "Exploring the Influence of Pore Shape on Conductance and Permeation"

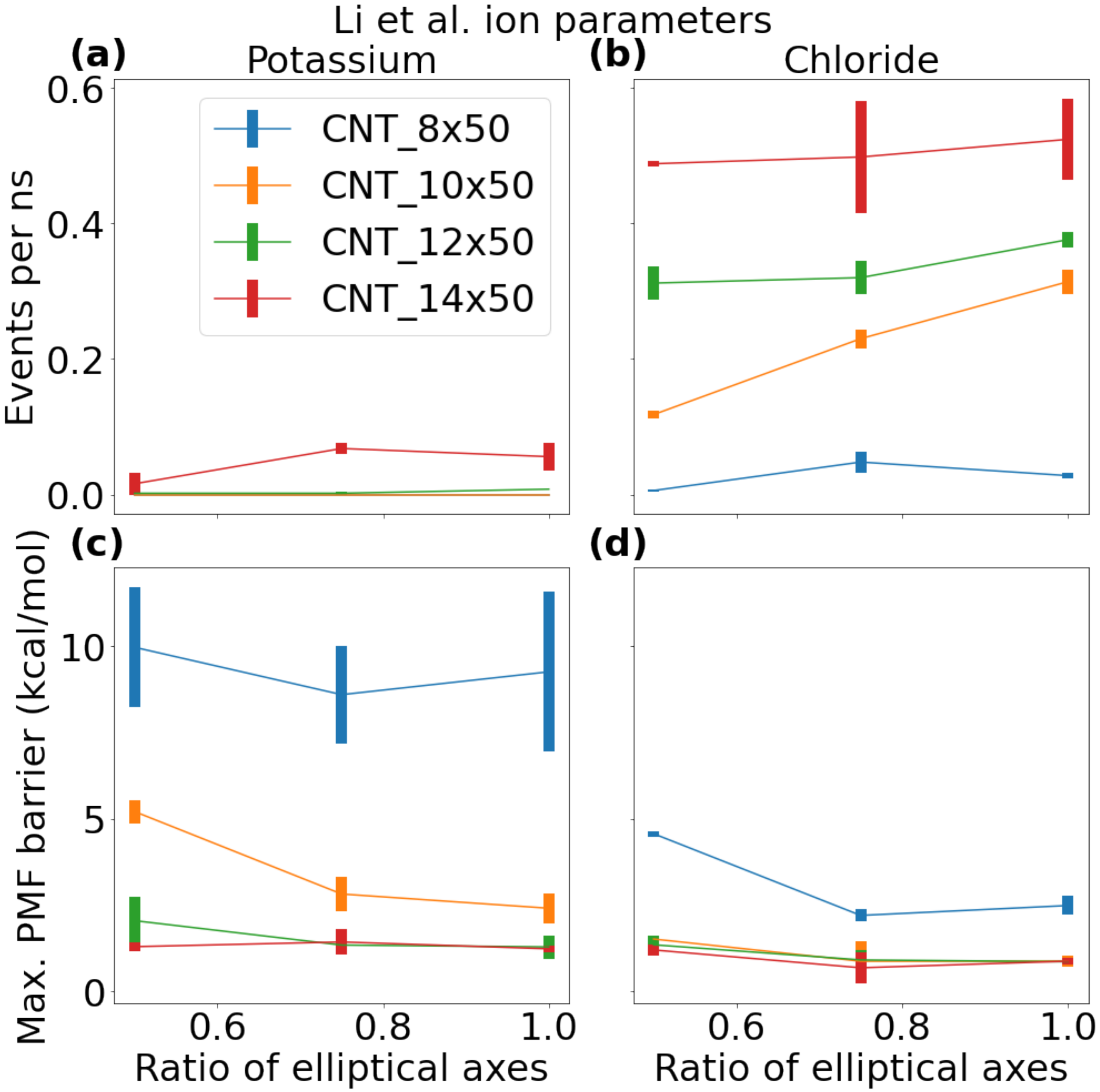


***Figure S1.*** *Barrier for ion permeation and ion conductance for carbon nanotubes (CNTs) as a function of the ratio between of the elliptic radii using ion parameters from Li et al (2015). The conductance is measured as events per ns observed in multiple 250 ns repeats with external potential of 500 mV for potassium* ***(a)*** *and chloride* ***(b)*** *ions . The energetic barrier is the maximum of the Potential of Mean Force (PMF) obtained from umbrella sampling and is computed for CNTs with potassium* ***(c)*** *and chloride* ***(d)*** *ions.*
